# TrimNN: Characterizing cellular community motifs for studying multicellular topological organization in complex tissues

**DOI:** 10.1101/2024.12.19.629384

**Authors:** Yang Yu, Shuang Wang, Jinpu Li, Meichen Yu, Kyle McCrocklin, Jing-Qiong Kang, Anjun Ma, Qin Ma, Dong Xu, Juexin Wang

**Author notes:** These Authors Contributed Equally (Yang Yu, Shuang Wang). Corresponding Authors: Qin Ma; Dong Xu; Juexin Wang.

## Abstract

The spatial organization of cells plays a pivotal role in shaping tissue functions and phenotypes in various biological systems and diseased microenvironments. However, the topological principles governing interactions among cell types within spatial patterns remain poorly understood. Here, we present the **Tri**angulation Cellular Community **M**otif **N**eural **N**etwork (**TrimNN**), a graph-based deep learning framework designed to identify conserved spatial cell organization patterns, termed Cellular Community (**CC**) motifs, from spatial transcriptomics and proteomics data. TrimNN employs a semi-divide-and-conquer approach to efficiently detect over-represented topological motifs of varying sizes in a triangulated space. By uncovering CC motifs, TrimNN reveals key associations between spatially distributed cell-type patterns and diverse phenotypes. These insights provide a foundation for understanding biological and disease mechanisms and offer potential biomarkers for diagnosis and therapeutic interventions.

## INTRODUCTION

Various cells work together within spatial arrangements in the tissue to support organ homeostasis and function^1^. Deciphering the multicellular organization is key to understanding the relationship between spatial structure and tissue biological and pathological functions^2^. Emerging spatial omics approaches, including spatially resolved transcriptomics^3^ and spatial proteomics^4^, enable investigation of the mechanisms governing the spatial organization of different cell types in a specific tissue. Within region of interest (ROI) in spatial omics, cellular neighborhoods (CNs) define local cell type enrichment patterns in cellular communities (CCs), and decoding function-related conservative spatial features in CNs is one of the primary spatial omics data analysis tasks^4^.

Most existing data analysis approaches adopt the top-down strategy to describe the cell organizations, which mainly relies on clustering strategies to identify the cell type compositions as common patterns. Deep learning approaches, including SPACE-GM^5^, CytoCommunity^6^, CellCharter^7^, and BANKSY^8^, typically learn low-dimensional embeddings of the nodes in corresponding CNs and then apply clustering approaches to these embeddings. However, clustering approaches suffer the following challenges in dissecting and interpreting highly heterogeneous, dynamically evolving cell systems^9^. First, clustering results usually become less stable when samples contain cells under active state transition, which is common in disease or developmental processes^10^. Second, clusters identified by these top-down approaches are often described as percentages of cell-type compositions. These clustering presentations lack formulations in topologically representing the geometrical cell-type interactions or are not easy to interpret biologically. Last, these top-down results essentially depend on the presence of batch effects, where CNs separate primarily by samples as technical covariates rather than biological features^3^. These batch effects make it easy to overfit the models but difficult to validate across different data sets^5^.

Considering the above limitations of top-down strategies, we identify CC motifs as recurring significant interconnections between cells using a bottom-up strategy. In the spatial omics-derived CC, we hypothesize that CC motifs can be represented as topological building blocks of multicellular organization consistent across different samples and associated with key biological processes and functions. CC motifs are biologically interpretable spatial patterns of the combined cell types, which provide topological information beyond clusters, and explicitly link to the biological and pathological mechanisms through distinct cell-cell communications, highly expressed genes and pathways^11^. This concept is related to the Functional Tissue Units (FTUs)^12^, but CC motifs are even smaller in the scale of cell locations and cell types, which provides more details for understanding and modeling the healthy physiological function of the organ and changes during disease states. Currently, size 1-3 motif analysis^4^ makes up most of the spatial omics studies, where motifs in size-1 as nodes can be treated as cell type compositions, size-2 as double nodes linked by edges, and size-3 as triple nodes within triangles. Nevertheless, biologists have found that sizable CC motifs with more nodes than triangles substantially correlate with patient survival and phenotypical features in colorectal cancer^13^, kidney diseases^14^, maternal-fetal interface^15^, and many more.

In practice, identifying the most overrepresented CC motifs composing multicellular organization is still computationally expensive with (*i*) subgraph matching^16^, which counts the occurrence of a given motif on the query graph, and (*ii*) pattern growth^17^, which finds the motifs with the most significant occurrence. It is known that subgraph matching is NP-complete^16^, which makes the node type combination alone super-exponential. Existing approaches include permutation^11^, edge sampling (e.g., MFinder^18^), node sampling (e.g., FANMOD^19^), and global pruning (e.g., Ullmann^20^ and VF2^21^). There is not yet a computationally feasible approach to analytically identify conservative, interpretable, and generalizable spatial rules of cellular organization in different sizes across different samples of spatial omics.

Here, we propose Triangulation cellular community Motif Neural Network (**TrimNN**), a graph-based deep learning approach to analyze spatial transcriptomics and proteomics data using a bottom-up strategy (**Supplementary Fig. 1**). Within the input spatial omics samples, CC is defined based on the cells as nodes, the node types represent different cell types, and the edges encode physical proximity inferred unidirectional as the spatial cell-cell relation from Delaunay triangulation^22^ based on nodes coordinates from ROI. TrimNN estimates overrepresented size-*K* CC motifs in the CC of spatial omics using graph isomorphism network^23^ (GIN) empowered by positional encoding^24^ (PE). In various spatial transcriptomics and spatial proteomics case studies, TrimNN identifies computationally significant and biologically meaningful CC motifs to differentiate patient survivals in colorectal cancer studies and represent pathological-related cell type organization in neurodegenerative diseases and colorectal carcinoma studies. Notably, the identified sizable CC motifs demonstrate their potential as interpretable topological prognostic biomarkers linking the topological structural organization of cell types at microscopic levels to phenotypes at macroscopic levels, which cannot be inferred by other existing tools. The source code of TrimNN is publicly available at https://github.com/yuyang-0825/TrimNN.

## RESULTS

### TrimNN quantifies multicellular organization with sizeable CC motifs

A schematic diagram of the proposed TrimNN and its analytic workflow is shown in **Fig. 1A**. These identified CC motifs are biologically interpretable through a set of downstream analyses, including motif visualization, cellular level interpretation within cell-cell communication analysis, gene level interpretation within differentially expressed gene and pathway analysis, and phenotypical analysis within the availability of phenotypical information (**Fig. 1B**). TrimNN is constructed on an empowered GIN to estimate the occurrence of the query on the target graph. On the CC as a triangulated graph built from spatial omics, it builds a supervised graph learning model by simplifying the graph constraints and incorporating the inductive bias within triangles from Delaunay triangulation. TrimNN decomposes the regression task in occurrence counting of the query graphs into many trackable binary classification tasks modeled by the sub-TrimNN module. Inspired by NSIC^25^, it is trained on representative pairs of the predefined query subgraphs and the target triangulated cell graphs as a binary classification task. This graph representation framework builds upon GIN and adopts a shortest-distance-based PE^24^, modeling the symmetric space to increase the expressive power. Additionally, TrimNN adopts a semi-divide and conquer strategy to estimate the abundance of the query by summarizing the enumeration of single classification tasks by sub-TrimNN module on each node’s enclosed graph. Given the size of the query subgraph, our framework uses an enumeration approach to estimate the most overrepresented CC motifs with possible cell types and topology. Then, we search to infer CC motifs in different sizes incrementally. The details of the architecture of TrimNN are shown in **Supplementary Fig. 2**.

**Figure 1.**
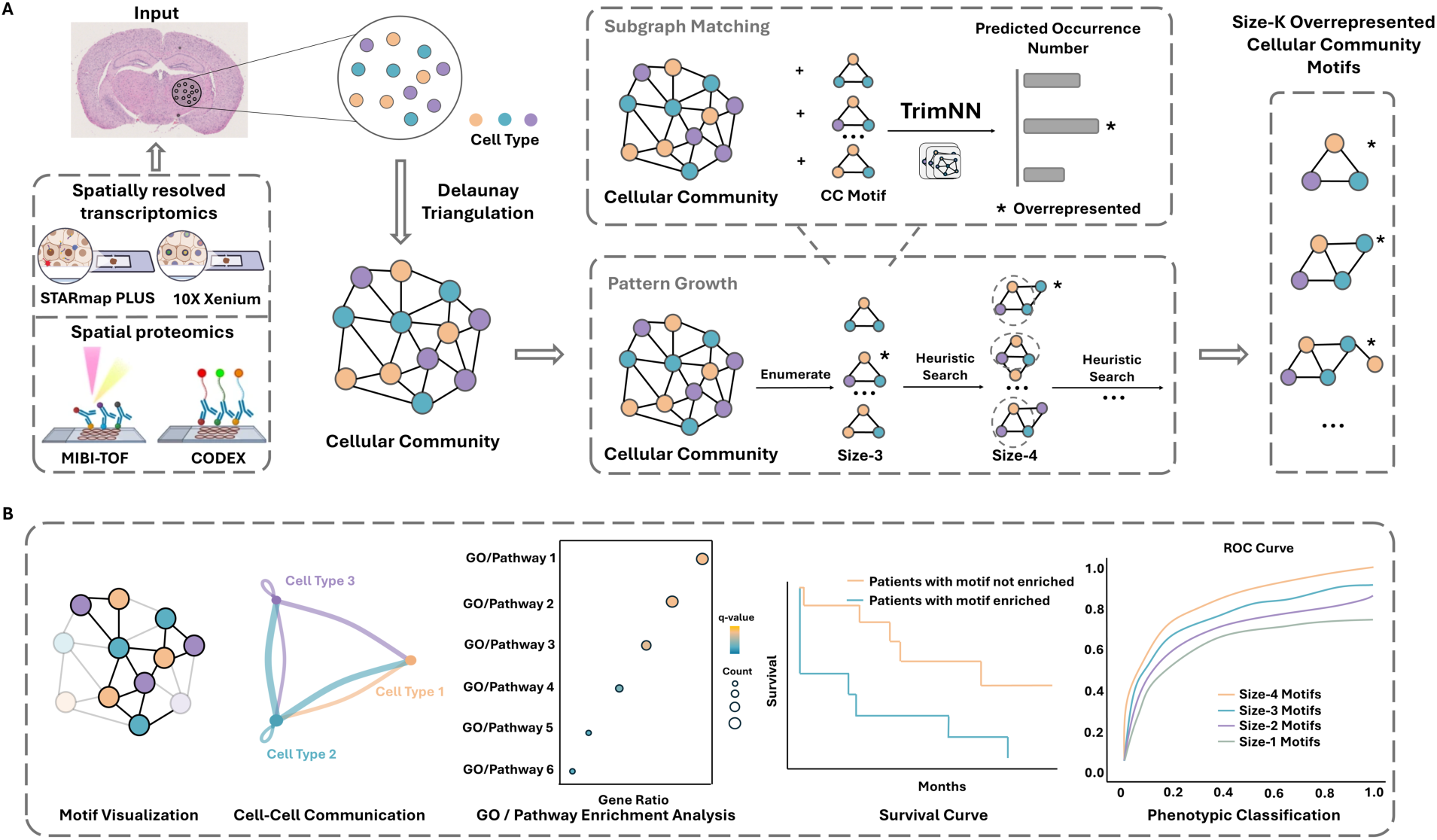
TrimNN analysis workflow. **A.** Spatially resolved transcriptomics, e.g., STARmap PLUS and 10X Xenium, and spatial proteomics data, e.g., MIBI-TOF and CODEX, are used as input to generate corresponding CC with spatial coordinates and Delaunay Triangulation. TrimNN is trained on representative pairs of query motifs and target triangulated graphs at scale. Given a specific query, TrimNN identifies its occurrence in the target CC in the subgraph matching process by decomposing this regression task to many binary classification problems, where each classification predicts whether the query exists in the target graph as the enclosed graph of each node. Enumerating possible motifs at size-*k*, TrimNN identifies the most overrepresented motifs. Then, the pattern growth process adopts a heuristic search for their successor size-*k* + 1 motifs. Here, we take size-3 CC motifs as an example. After subgraph matching and pattern growth, TrimNN estimates overrepresented CC motifs. **B.** These CC motifs can be biologically interpreted in the downstream analysis, including visualization, cellular level interpretation within cell-cell communication analysis, gene level interpretation within differentially expressed gene analysis, e.g., GO enrichment analysis and pathway enrichment analysis, and phenotypical analysis within the availability of phenotypical information, e.g., survival curve and phenotypic classification analysis. CC: cellular community.

We hypothesize CC motifs as the countable recurring spatial patterns of various cell types are robust within noises to represent and quantify multicellular organization. Simulations were performed to mimic different levels of noises, including cell missing from the cell capture imperfection of sequencing technology (**Fig. 2A**), cell coordinates shifting from technological errors (**Fig. 2B**), and cell type misclassification from annotation errors in data analytics (**Fig. 2C**). We noticed diverse noises do not influence the relative ranking of CC motif abundance, they were robustly consistent in most scenarios (**Fig. 2A** and **Supplementary Fig. 3**). Even under extreme cases with a noise ratio of 0.5, the Spearman correlation in cell missing and cell coordinates shifting was above 0.92 and 0.97, respectively, while errors from cell type annotation still made the CC motif abundance manageable around 0.75.

**Figure 2.**
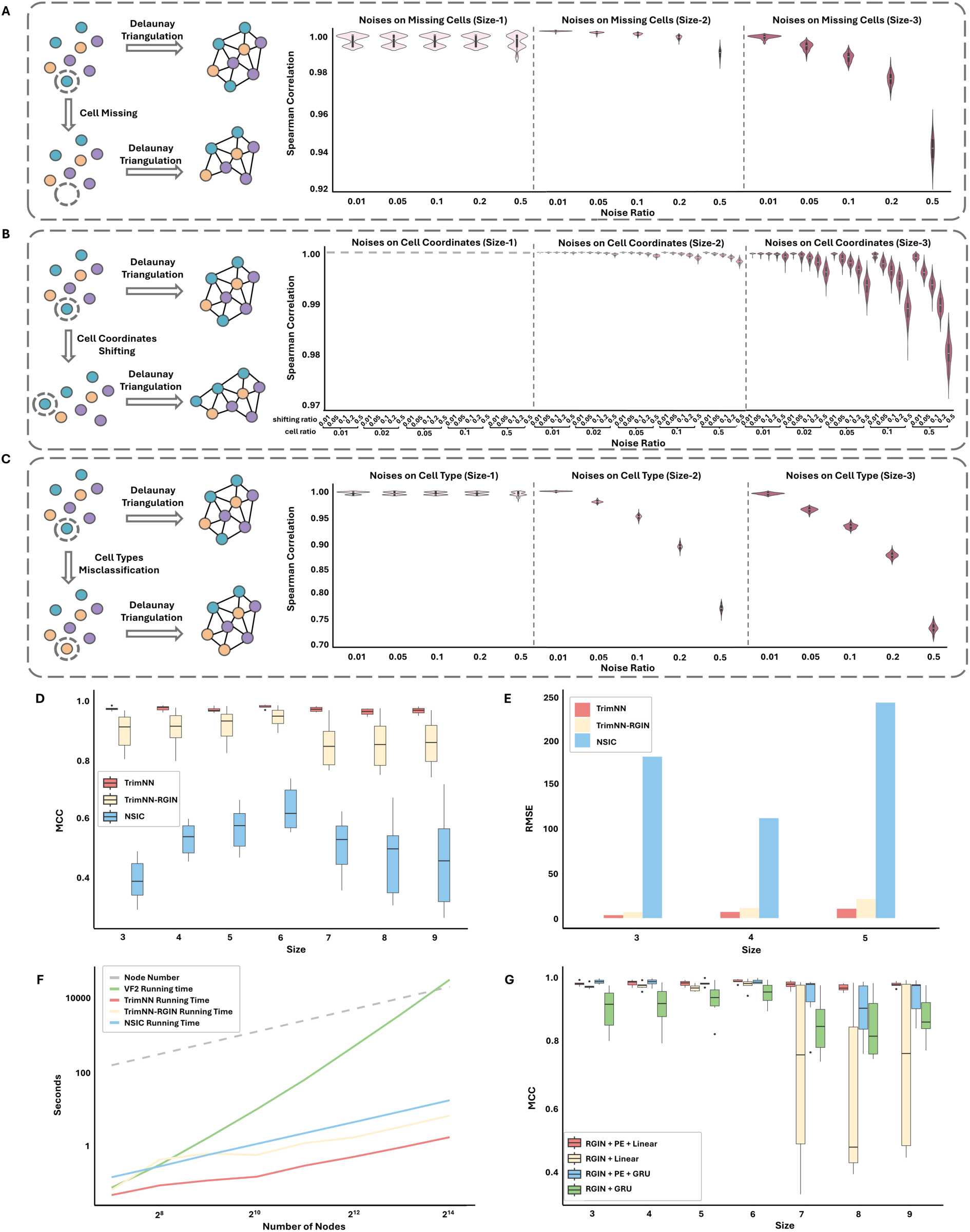
The performance of TrimNN on spatial omics. **A**. Simulations of cell missing effects on CC motifs, represented as Spearman correlation of rankings of abundances before and after simulated noises at cell proportions of 0.01, 0.05, 0.1, 0.2, and 0.5 within CC motifs in size-1, size-2, and size-3. **B**. Simulations of cell coordinate shifting effects on CC motifs, represented as Spearman correlation of rankings of abundances before and after simulated noises with different levels of noises of 0.01, 0.05, 0.1, 0.2, and 0.5 at cell proportions of 0.01, 0.05, 0.1, 0.2, and 0.5 within CC motifs in size-1, size-2, and size-3. **C**. Simulations of cell-type misclassification effects on CC motifs, represented as Spearman correlation of rankings of abundances before and after simulated noises at cell proportions of 0.01, 0.05, 0.1, 0.2, and 0.5 within CC motifs in size-1, size-2, and size-3. **D.** Benchmarking the performance of TrimNN, TrimNN-RGIN, and NSIC on independent simulated data for subgraph matching. The X-axis represents different sizes of CC motifs, and the Y-axis indicates the MCC (Matthews Correlation Coefficient) values. **E.** Performance comparison of TrimNN, TrimNN-RGIN, and NSIC in identifying occurrences of CC motifs in diverse simulated data sets. The Y-axis is the RMSE (Root Mean Square Error) value. **F.** Scalability of TrimNN. The X-axis represents the size of the triangulated graph, and the Y-axis indicates the runtime on a workstation equipped with an Intel Xeon Gold 6338 CPU and 80G RAM. **G.** Ablation tests on performance comparison adding positional encoding of TrimNN model. The X-axis represents different sizes of CC motifs, and the Y-axis indicates the MCC values. CC: cellular community.

### TrimNN accurately identifies overrepresented CC motifs in Cellular Neighborhoods

On a modified subgraph matching task as a binary classification of motif existences in a triangulated graph, TrimNN outperformed the competitive methods in all scenarios in most criteria in synthetic spatial omics data, including VF2, original regression-based neural network method NSIC^25^, and TrimNN-RGIN with proposed formulation but using NSIC’s RGIN network architecture. Especially on large-size CC motifs, TrimNN demonstrated significant performance improvements with TrimNN-RGIN, highlighting its architecture’s capacity (**Fig. 2D** and **Supplementary Data 1**).

TrimNN accurately identified the top overrepresented CC motifs. On a pattern growth challenge to determine the ranking of CC motifs abundance, TrimNN was shown to outperform the competitive methods consistently in different sizes and cell types in synthetic spatial omics data. Both TrimNN and TrimNN-RGIN outperformed NSIC by a large margin in most scenarios and criteria, which highlights the capability of the proposed problem formulation. Notably, TrimNN demonstrated an average improvement over NSIC by approximately 20∼60 times in Root Mean Square Error (RMSE) (**Fig. 2E** and **Supplementary Data 2**). Besides criteria in absolute occurrence value, the relative value of the ranking index also supported TrimNN’s capacity in **Supplementary Fig. 4A** and **Supplementary Data 3**.

TrimNN is highly scalable in identifying large-size CC motifs. As scalability plays a vital role in the study, we compared the computational time on target triangulated graphs with varying node sizes. We observed TrimNN, TrimNN-RGIN, and NSIC exhibit linear scalability with increasing node sizes, while TrimNN continuously consumed lower computational time (**Fig. 2F**). Especially, TrimNN was more efficient than TrimNN-RGIN with a simplifier network architecture using same problem setting. In contrast, the classical enumeration-based VF2 method grew exponentially, where its runtime made it unacceptable in most scenarios. In practical usage, on typical spatial omics data with thousands of cells of dozens of cell types, TrimNN robustly infers large-size CC motifs accurately in seconds, which is unattainable through conventional methods.

PE together with GIN increases the expressive power of TrimNN. In challenging tasks with larger-sized motifs, ablation tests showed that integrating PE improved GNN performance compared with TrimNN-RGIN without PE and a complex GRU module (**Fig. 2G** and **Supplementary Data 4**). In addition, GIN as the critical component in TrimNN was effective by replacing it with other graph neural network models including Graph Convolutional Networks and Graph Transformer^26^, keeping other components and parameters constant (**Supplementary Fig. 4B** and **Supplementary Data 5**). This result aligned with theoretical analyses that GIN is a powerful 1-order graph neural networks^23^. Meanwhile, it was shown that TrimNN requires sufficient training data to learn the complex relationships (**Supplementary Fig. 4C** and **Supplementary Data 6**).

### TrimNN identifies representative CC motifs that accurately differentiate the severity of colorectal cancer patients

In addition to the above simulation studies, we showed that the CC motifs inferred by TrimNN are intrinsic representations to differentiate phenotypes of the CC. In a proteomics study using Co-Detection by Indexing (CODEX)^13^ on colorectal cancer (CRC) that contains 17 low-risk (Crohn’s-like lymphoid reaction (CLR)) and 18 high-risk (diffuse inflammatory infiltration (DII)) patients of 140 tissue regions, we performed a CC motif analysis using TrimNN and identify most abundant CC motifs in sizes 1-4. Traditional machine learning approaches such as Logistics Regression (LR) were adopted using relative ranking indices to quantify motif occurrence as features. As the original publication annotated 29 cell types, we chose 29 as the fixed number of features in supervised learning to classify CLR and DII. Within 10-fold cross-validation following the same protocol as CytoCommunity^6^, the ROC-AUC results of LR were 0.77, 0.76, 0.79, and 0.76 (**Fig. 3A**) for size 1-4 CC motifs, respectively. This LR model with 29 CC motif features outperformed CytoCommunity’s extensive GNN computation performance using 512 dimensions of embeddings as features (ROC-AUC: 0.71). Notably, if the feature number increased to the top 100, the LR model on size-3 motifs achieved an ROC-AUC of 0.81. In addition, other classical machine learning models, such as Radom Forest and Support Vector Machine, were applied to the same classification tasks with the same settings. These models performed similarly to LR, further supporting the representational power of CC motifs (**Supplementary Data 7-9**).

**Figure 3.**
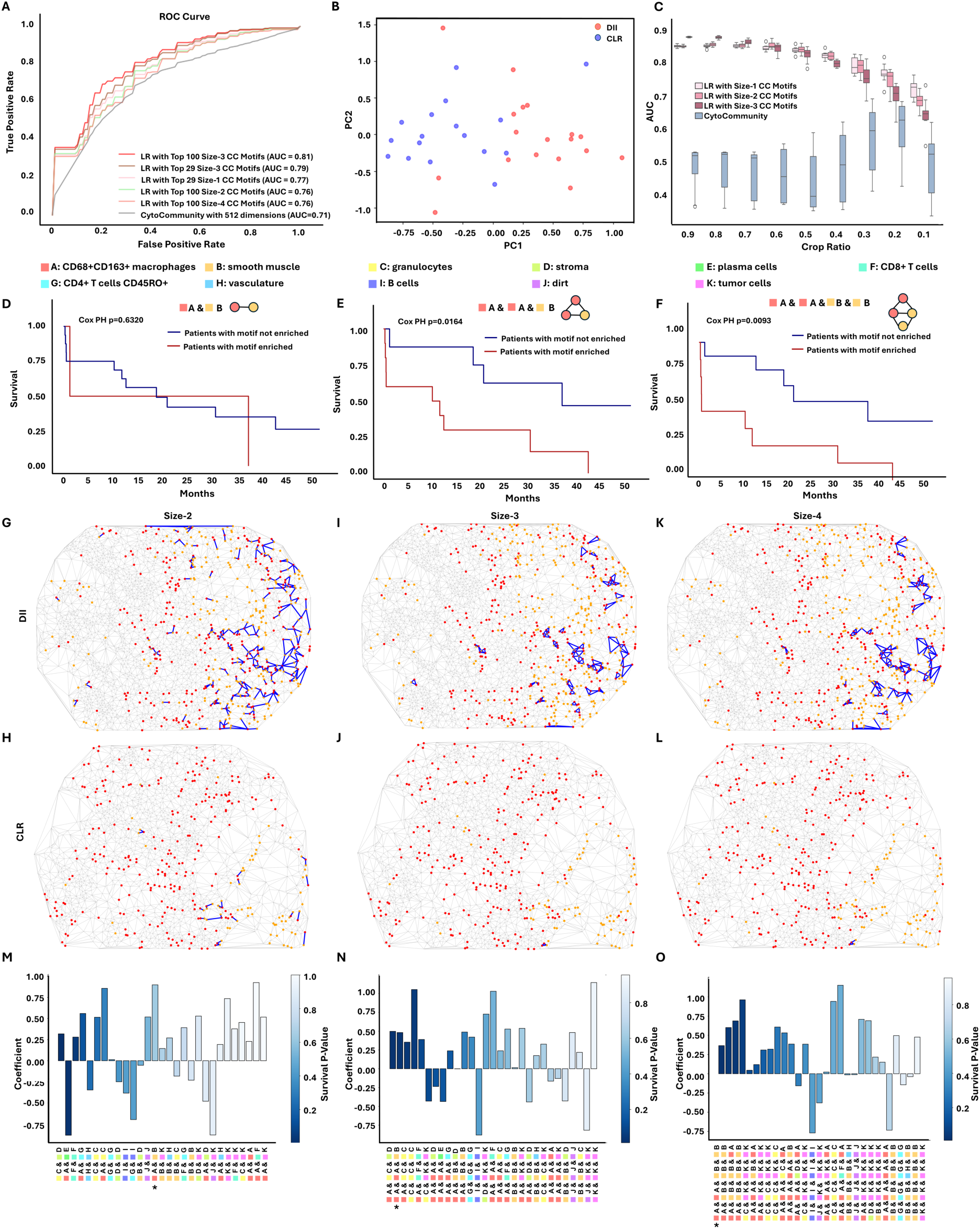
TrimNN analysis on a colorectal cancer study using CODEX. **A**. The ROC curves of the Logistics Regression model classify CLR and DII patients using top CC motifs of size 1-4 as features, and the competitive method CytoCommunity uses learned dimension. The Logistic Regression model features motif counts from TrimNN and scales between 0 and 1. **B**. Visualize all samples using the top two principal components from 29 top size-2 CC motifs. Blue spots denote the CLR patient group, and red spots denote the DII patient group. **C.** Generalizability of the trained model testing on random cropping of ROI in the samples. X-axis is the ratio of width and height of the original ROI, Y-axis is ROC-AUC. Survival curves of DII patients with and without enriched motifs, including **D**. size-2 ‘A & B’. Here cell type CD68+CD163+ macrophages are denoted as ‘A’, and smooth muscles are denoted as ‘B’, **E**. size-3 ‘A & A & B’, and **F**. size-4 ‘A & A & B & B’. The visualization of spatial localization of size-2 CC motif ‘A & B’ on the CC in **G**. patient 3 (DII) on spot 5A and **H**. patient 8 (CLR) on spot 16A. The visualization of spatial locations of the size-3 motif ‘A & A & B’ in **I.** DII spot and **J.** CLR spot (same spots as **G** and **H**). The visualization of spatial localization of the size-4 motif ‘A & A & B & B’ in **K.** DII spot and **L.** CLR spot (same spots as **G.** and **H.**). All motifs are marked as blue, nodes of cell type A are red, and nodes of cell type B are orange. The plot of Logistic Regression coefficients ranked by Cox PH value of the top 29 CC motifs in **M**. size-2, **N.** size-3, and **O.** size-4. The extent of the blue color represents the Cox PH value. * marks the highlighted motif. CC: cellular community.

Besides supervised learning, these CC motifs seemed to capture some intrinsic characteristics in CNs, where CLR and DII demonstrated good visual separation using the top two principal components inferred from the top 29 size-2 motifs (**Fig. 3B**). Further unsupervised hierarchical clustering showed the different distribution of top 29 motif abundance among CLR and DII groups in **Supplementary Fig. 5A, 5B** (size-2), **Supplementary Fig. 5C, 5D** (size-3), and **Supplementary Fig. 5E, 5F** (size-4).

CC motifs in simpler models and fewer numbers of features showed better generalizability across multiple samples in machine learning. When only parts of the CC were available by random cropping the samples, CC motifs-based LR methods were very robust in generalization compared to competitive methods (**Fig. 3C**). The same trends were observed in distorted samples with simulated noises in cell missing, cell coordinates shifting, and cell type misclassification (**Supplementary Fig. 6**).

In addition, the enrichment of sizable CC motifs can be used to differentiate patient survival. We identified several size-2, 3, and 4 CC motifs that significantly differentiate survival (Cox PH < 0.05) between enriched and non-enriched DII patients, while cell type composition (size-1 motifs) may not necessarily succeed (**Supplementary Data 10**). With cell type ‘CD68+CD163+ macrophages’ (denoted by ‘A’) and cell type ‘smooth muscle’ (denoted by ‘‘B’), the survival curves showed that size-2 motif ‘A & B’ may not have well-separated survival in the DII patient group (Cox PH = 0.63, shown in **Fig. 3D**), but including more adjacent nodes with the same cell types, patients with enrichment of size-3 ‘A & A & B’ and size-4 CC motifs ‘A & A & B & B’ showed significant lower survival rate (Cox PH = 0.016, shown in **Fig. 3E**, and Cox PH = 0.0093, shown in **Fig. 3F**, respectively). In addition, the occurrence numbers of these CC motifs among DII and CLR patients were 14,415 and 7,004 (ratio 2.06) for size-2 ‘A & B’, 4,176 and 1,548 (ratio 2.70) for size-3 ‘A & A & B’, 6,946 and 2,276 (ratio 3.05) for size-4 ‘A & A & B & B’, all were inferred as significance through Benjamini-Hochberg adjusted Fisher’s exact test. The different distribution of these CC motifs on CCs among DII and CLR spots can be visualized in **Fig. 3G, 3H.** (size-2), **Fig. 3I, 3J.** (size-3), and **Fig. 3K, 3L.** (size-4).

Notably, spatial topology plays a crucial role in linking phenotypes and survival. There were two types of size-3 motifs with cell types CD68+CD163+macrophages (‘A’) and smooth muscle (‘B’). Compared with ‘A & A & B’, the alternative motif ‘A & B & B’ had occurred 4,602 and 2,255 times among DII and CLR patients with a lower ratio of 2.04, and it cannot differentiate survival well (Cox PH = 0.2975). Apparently, these topological differences among spatial localization of cells in different cell types played different roles biologically and pathologically, where conventional top-down approaches with cell type composition failed to distinguish (**Supplementary Fig. 7** and **Supplementary Data 11-14**).

Furthermore, an LR model provides intrinsic interpretability when differentiating phenotypes. The coefficients of each feature from the LR model demonstrated CC motifs’ importance quantitatively, making the model interpretable (**Fig. 3M**, **3N**, and **3O**). Notably, all macrophage and muscle-related and significant Cox PH motifs in different sizes tended to have high absolute coefficient values. The same interpretable results can also be cross-validated by Shapley value^27^ in **Supplementary Fig. 5G, 5H**, and **5I**, showing these macrophage and muscle-related CC motifs were essential to differentiating DII patients from CLR patients. Biologically, it was evidenced that macrophages facilitate pancreatic cancer to induce muscle wasting via promoting TWEAK (TNF-like weak inducer of apoptosis) secretion from the tumor^28^. After carefully checking these top motifs in different sizes, we also identified biologically meaningful tumor cells and B cells, which were known to be related to the severity of CRC^29^. Representative tumor and B cell enrichments in DII and CLR samples were shown in **Supplementary Fig. 5J, 5K, 5L**, and **5M**. Our analysis validated the crosstalk between macrophages, muscle wasting, and cancer cachexia through an independent spatial omics study, and TrimNN identified CC motifs in a data-driven approach as robust interpretable representations in CNs.

### TrimNN identifies CC motifs revealing diverse roles in Alzheimer’s disease using spatial transcriptomics data

Next, we showed TrimNN’s capability to identify diverse spatially distributed CC motifs corresponding to multiple biological and pathological mechanisms in complex diseases. It is known that the interaction between cortical excitatory neurons and microglia is significantly disrupted by neuroinflammation in AD^30^. However, their topological combinations, particularly their relationship with amyloid-β on the cellular level, are still unknown^31^. We performed TrimNN on an Alzheimer’s disease (AD) mouse brain study with eight-month-old and thirteen-month-old samples sequenced by STARmap PLUS spatially resolved transcriptomics^32^. There were two replicates for both disease and control conditions at each time point. The transcriptomics data included 2,766 genes and two proteomics channels representing AD markers of amyloid-β and tau pathologies at subcellular resolution.

On the derived CCs, size-3 triangle-like CC motifs composed by cell types cortex excitatory neuron and microglia were identified significant between AD (**Fig. 4A**) and control (**Fig. 4B**). These significant CC motifs included Cortex-Cortex-Cortex (CCC), Cortex-Cortex-Microglia (CCM), Cortex-Microglia-Microglia (CMM), and Microglia-Microglia-Microglia (MMM) (Benjamini-Hochberg adjusted Fisher’s exact test in eight-month-old replicate 1 with *p-values* 6.12e-32, 3.23e-20, 4.30e-34, and 9.74e-07, respectively). Visualization of an exemplary CC motif ‘MMM’ demonstrated uneven spatial distribution that differed in AD (**Fig. 4C**) and control (**Fig. 4D**). **Supplementary Fig. 8** showed the spatial occurrence distribution of the other three motifs. The CCs inferred from all samples were shown in **Supplementary Fig. 9** and **Supplementary Data 15-26**.

**Figure 4.**
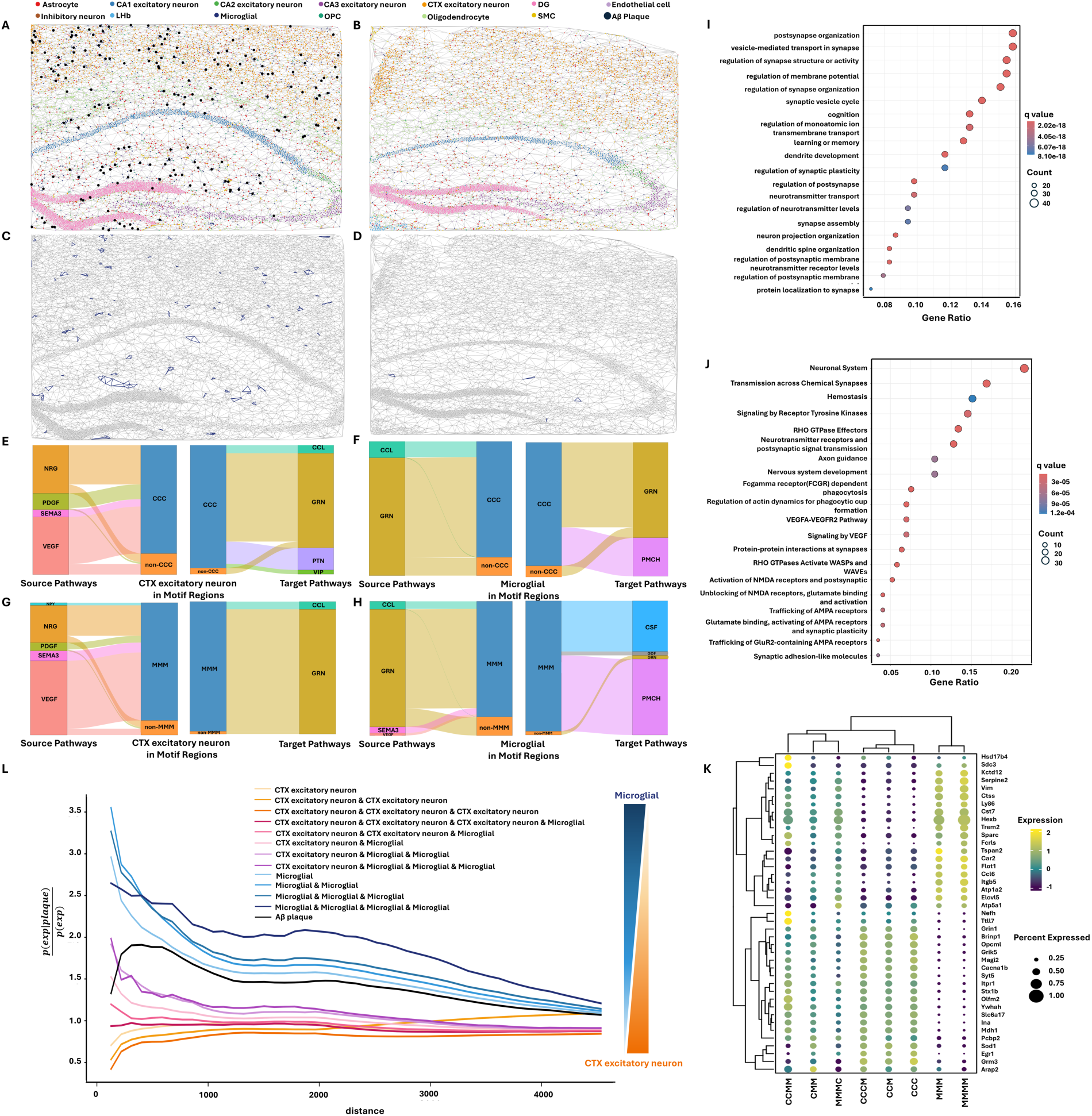
TrimNN analysis on an AD mouse study sequenced by STARmap PLUS. CCs of thirteen-month-old of **A.** AD and **B.** control sample replicate 1 using Delaunay Triangulation, where black spots are amyloid-β in the AD sample. The spatial locations of the identified motif with all microglia cells (‘MMM’ motif, where ‘M’ denotes cell type microglia) in thirteen-month-old replicate 1 of **C.** AD and **D.** control mouse samples. ‘MMM’ motifs are marked as purple. Cell-cell communication analysis demonstrates the ligand-receptor differences between motif regions and non-motif regions as river plots, including **E.** cell type cortex excitatory neurons (denoted as ‘C’) as source (left) and as target (right), **F.** cell type microglia as source (left) and target (right) in regions with and without ‘CCC’ motif. Similarly, **G.** and **H.** are cell-type Cortex excitatory neurons and Microglia as source and target in regions with and without the ‘MMM’ motif. **I.** GO enrichment analysis of Biological Processes and **J.** Pathway enrichment analysis on DEGs between regions containing and not containing the ‘MMM’ motif in thirteen-month-old AD samples. **K.** Expression of marker genes for cell type ‘C’ and ‘M’ related size-3, size-4 motifs. **L.** Spatial co-occurrence of different CC motifs with respect to amyloid-β as computed using Squidpy. Microglia-related motifs have even higher spatial co-occurrence probability to the amyloid-β plaque, and cortex excitatory neuron-related motifs have lower spatial co-occurrence probabilities to the amyloid-β plaque. CC: cellular community. CTX: Cortex.

Here, we defined motif-enriched regions as expanded regions within three hops of CC motifs in the CC. From the perspective of cell-cell communications, unique ligand-receptor signaling pathway patterns between ‘CCC’ (**Fig. 4E-F**) and ‘MMM’ (**Fig. 4G-H**) were identified by motif-enriched and complementary regions on thirteen-month-old samples using CellChat^33^. Specific to cell type microglia, motifs ‘CCC’ and ‘MMM’ had dominant ligand-receptor pairs *GRN*^21^ and *PMCH* to distinguish motif regions from the complementary regions. AD-related ligand-receptors, including *GRN*, *VEGF*, *PDGF*, *CCL*, *VIP*, *NRG*, and *SEMA3*, were significantly enriched in CC motifs associated with cortex excitatory neurons or microglia (**Supplementary Data 27-30**). All the cell-cell communication results on CC motifs were detailed in **Supplementary Figs. 10-18** and **Supplementary Data 31-38**.

Then we analyzed the gene-level characteristics of identified CC motifs. Comparing ‘MMM’ motif-enriched and complementary regions, differentially expressed genes (DEGs) were identified as significant (*p*-value<0.05) using DESeq2^34^, including *Plekha1*, *Ctsb*, *Sort1* in eight-month-old samples (**Supplementary Data 42**), and *App*, *Plekha1*, *Clu*, *Ptk2b*, *Sort1, Bin1*, *Ctsb* in thirteen-month-old samples(**Supplementary Data 46**). On DEGs in thirteen-month-old samples, Gene Ontology (GO) enrichment analysis showed significant vesicle-mediated transport in synapse (*q*-value 5.90e-36), regulation of synapse structure or activity (*q*-value 5.67e-32), learning or memory (*q*-value 4.68e-24), and cognition (*q*-value 1.40e-22) (**Fig. 4I**). Neural systems (*q*-value 8.32e-16), Transmission across Chemical Synapses (*q*-value 6.30e-15), Neurotransmitter receptors and postsynaptic signal transmission (*q*-value 4.84e-12), and Nervous system development (*q*-value 6.83e-05) were enriched with pathway enrichment analysis (**Fig. 4J**). For all the detailed results of CC motifs ‘CCC’, ‘CCM’, ‘CMM’, and ‘MMM’, a similar analysis was performed for DEGs (**Supplementary Data 39-46**), including GO enrichment analysis and pathway enrichment analysis (**Supplementary Figs. 19-20**).

To validate their relations with AD, we compared these motifs-related DEGs with 77 AD-associated genes identified from large-scale GWAS analysis^35^. On CC motif ‘CCC’, the *Trem2* gene was exclusively observed in the replicates of the thirteen-month-old but not eight-month-old AD mouse model, consistent with its role in the late-onset of AD^36^. Similarly, for the motif ‘MMM’, the *Clu* gene was highlighted only in the thirteen-month-old mouse model, aligning with its direct involvement in the formation process of amyloid-β^37^ (**Supplementary Fig. 21**).

Based on the identified size-3 motifs, we performed pattern growth to identify size-4 motifs using TrimNN (**Supplementary Notes 1**). Among all the size-4 ‘CCC’ expanded motifs, ‘CCCM’ showed the most significant difference between AD and control samples, while ‘MMMM’ was the most significant size-4 motif expanded from ‘MMM’. Similar to the analysis on size-3 motifs, checking significantly enriched ligand-receptors (**Supplementary Data 47-49**) and DEGs (**Supplementary Data 50-55**), these size-4 motifs were related to AD in cell-cell communication analysis (**Supplementary Data 56-61**), GO enrichment analysis (**Supplementary Fig. 22**), and pathway enrichment analysis (**Supplementary Fig. 23**).

Further investigation on DEGs showed diverse groups of CC motifs with expressed markers (**Fig. 4K**). Size-3 ‘MMM’ and size-4 ‘MMMM’ motifs with homogenous microglia had divergent expression patterns. For example, *Hexb* had a higher average expression than the other CC motifs. *Hexb* is known to induce toxic and progressive neuronal damage, which may relate to neurodegenerative dementia^38^.

In addition to examining the diversity of CC motifs at the gene level, we investigated whether the identified CC motifs were spatially colocalized with amyloid-β by computing their co-occurrence probabilities using Squidpy^39^. The results showed that microglia-related CC motifs had an even higher co-occurrence probability with amyloid-β than the spatial expectation, distinguishing them from other CC motifs associated with cortex excitatory neurons (**Fig. 4L**). Interestingly, the extent of homogeneity of microglia regions seemed to correspond to a larger co-occurrence probability to amyloid-β. In contrast, the extent of homogeneity of cortex excitatory neurons tallied to lower co-occurrence probability. This trend prevailed from the whole spectra of CC motifs composed of microglia and cortex excitatory neurons in multiple sizes, from a very high ratio of size-4 ‘MMM’ to a very low ratio of size-3 ‘CCC’.

Both differences in DEGs and spatial co-occurrence suggest the presence of two distinct types of CC motifs related to amyloid-β in AD. One type of CC motif, i.e., ‘CCC’, ‘CCM’, ‘CMM’, and ‘CCCM’, were reluctant to colocalize amyloid-β colocalization. Another kind of CC motif, i.e., ‘MMM’ and ‘MMMM’, were closely colocalized with amyloid-β. These results were consistent with the observation that microglia, as key mediators in the brain, activate inflammation in the vicinity of amyloid-β deposits, which are directly toxic to the adjacent neurons^40^. Activated microglia release pro-inflammatory cytokines, such as tumor necrosis factor-alpha (TNF-α) and interleukin-1 beta (IL-1β), which can damage excitatory neurons or alter their function^41^. In this case study, TrimNN facilitated the analysis of the spatial characteristics of microglia and cortex excitatory neurons, along with their topological relations with diverse cell types. TrimNN accurately captured their spatial colocalization patterns with amyloid-β deposits, providing insights into the onset of AD as the result of interactions between multiple cell types^42^.

### TrimNN identifies cell-type-specific spatial tendencies in colorectal carcinoma study on spatial proteomics data

Besides the AD study, we also performed TrimNN analysis to explore cell-type-specific spatial tendencies on one colorectal carcinoma study. It is known that the tumor microenvironment can significantly influence the interactions between T-cells and epithelial cells through antigen presentation, T-cell activation, and modulation of the tumor microenvironment. However, it is still unknown how the spatial arrangement of these cells is related to effective immune surveillance and the potential for therapeutic interventions^43^. The adopted colorectal carcinoma study investigated 40 ROIs in two colorectal cancer patients and 18 ROIs in two healthy controls using spatial proteomics of multiplexed ion beam imaging using time of flight (MIBI-TOF)^44^.

After a comprehensive analysis of size-3 and their related size-4 CC motifs with TrimNN, we defined two types of CC motifs: Shifted Interaction Motifs and Homeostatic Interaction Motifs (**Fig. 5A**). Shifted Interaction Motif demonstrated a shift of CC motif abundance from control-enrich (more occurrence in control than disease samples) to disease-enrich (more occurrence in disease than control samples) when expanding from size 3 to size 4. The exemplary size-3 motif ‘ABC’ (**Fig. 5B** and **5C**) suggested disease progression when involving other Immune cells to form a size-4 motif ‘ABCD’ (**Fig. 5D** and **5E**), where ‘A’ denotes CD4 T-cells, ‘B’ denotes CD8 T-cells, ‘C’ denotes Epithelial, and ‘D’ denotes other immune cells (other CD45+) annotated by the original publication. Proportion tests showed that this size-4 ‘ABCD’ significantly differed from the size-3 ‘ABC’ motif in abundance between disease and control samples (*p-*value 2.58e-12). In contrast, Homeostatic Interaction Motif remained consistent in abundance between disease and control groups when expanding its sizes, e.g., size-3 motif ‘AEC’ (**Fig. 5F** and **5G**), where ‘E’ denotes Endothelial concatenating another Epithelial(C) as a size-4 motif ‘AECC’ (**Fig. 5H** and **5I**). Proportion tests showed consistency between disease and control ratios among this pair of size-3 and size-4 motifs (*p-*value 0.55). The expression level of antibodies also confirmed differences between these two groups of CC motifs (**Fig. 5J, Supplementary Fig. 24A and 24B**). A similar analysis demonstrated that these two groups of CC motifs also existed in AD studies (**Supplementary Notes 2**).

**Figure 5.**
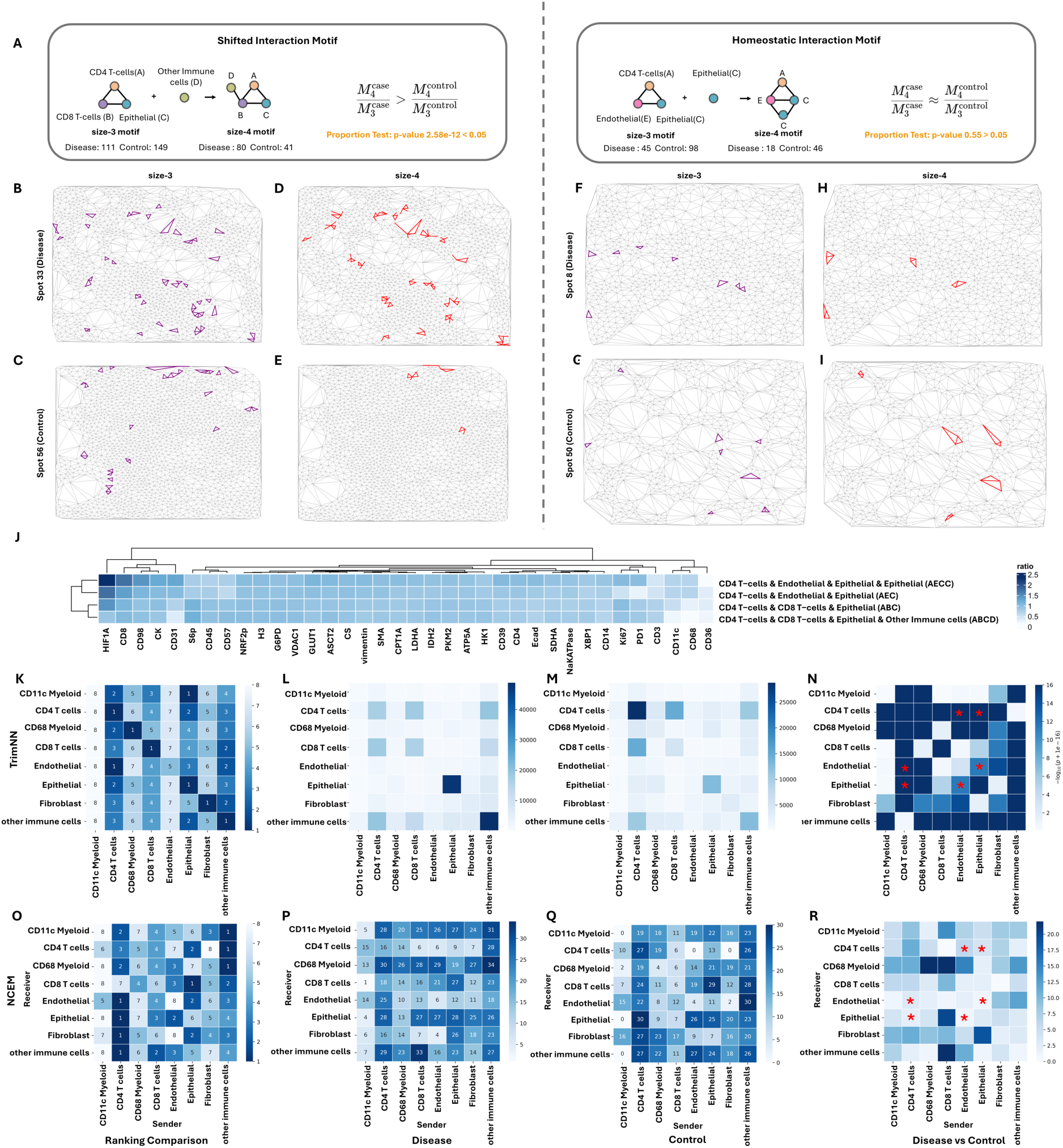
TrimNN analysis on a colorectal carcinoma study using MIBI-TOF. **A.** Schematic of Shifted Interaction Motif and Homeostatic Interaction Motif as two types of size-4 motifs. Shifted Interaction Motifs: size-3 motif ‘ABC’ (purple) in exemplary **B.** spot 33 (Disease) and **C.** spot 56 (Control), the successor size-4 motif ‘ABCD’ (red) in the same **D.** spot 33 (Disease) and **E.** spot 56 (Control). Homeostatic Interaction Motifs: exemplary size-3 motif ‘AEC’ (purple) in exemplary **F.** spot 8 (Disease) and **G.** spot 50 (Control), the successor size-4 motif ‘AECC’ (red) in the same **H.** spot 8 (Disease) and **I.** spot 50 (Control). **J.** Heatmap of antibody expression ratio between disease and control samples in Shifted Interaction Motif and Homeostatic Interaction Motif. **K**. Ranking of effective size between all cell types in colon tissue samples. Abundance of size-2 CC motifs as occurrences in **L**. Colon Carcinoma and **M**. Healthy control samples. **N**. P-value of size-2 CC motifs between disease and control by Benjamini-Hochberg adjusted Fisher’s exact test. **O**. Heatmap on sender rank from NCEM type coupling analysis in colon tissue samples. Heatmap of NCEM type-coupling analysis in **P**. Colon Carcinoma and **Q**. Healthy control samples. **R**. Difference values from NCEM type-coupling analysis between disease and control samples. Cell type ‘A’ denotes CD4 T-cells, ‘B’ denotes CD8 T-cells, ‘C’ denotes Epithelial, ‘D’ denotes other immune cells annotated by the original publication, and ‘E’ denotes Endothelial. The star symbol marks the paired cell type composition of the ‘AEC’ motif. CC: cellular community.

Next, we explored cell-type preferences using TrimNN analysis and identified CD4 T-cells that played key spatial roles in differentiating patients and healthy controls. Scrutinizing spatial colocalizations between cells in different cell types, homogeneous CD4 T-cells were most abundant in healthy control samples but ranked fifth in disease samples (purple rectangle in **Fig. 5K**) by effective sizes, with the *p-*value<10e-30 using Benjamini-Hochberg adjusted Fisher’s test (**Fig. 5L, 5M and Supplementary Data 62**). This result was also supported by differences in the sender-receiver effect by NCEM^45^ which estimates cell interactions of the effects of niche composition (**Fig. 5O, 5P**, and **5Q**). In addition, CD4 T-cells and epithelial cells tended to be more likely to be localized together than other combinations (yellow rectangle in **Fig. 5K**). This combination was also significantly differentiated between disease and control (adjusted *p-*value<10e-30), supported by the observation that epithelial as receiver significantly differs antibody expression in CD4 T-cells sender effect (**Supplementary Fig. 24C** and **24D**) from NCEM. Statistical coupling analysis within all the cell types between disease and control samples (**Fig. 5N**) showed that many were consistent with NCEM type coupling analysis (**Fig. 5R**). All these observations were consistent with the original studies^44^.

We also investigated the motifs with sizes larger than two where NCEM cannot inferred directly. The abundance of CC motif ‘AEC’ (‘CD4 T cells & Endothelial & Epithelial’) was observed to be significantly different (*p-*value 4.61e-15) between disease and control (**Supplementary Data 63**), where their composed pairs of cell types as size-2 motifs were confirmed as significant by TrimNN (**Fig. 5N**). However, NCEM failed to support this observation, for none of these pairs were shown to be significantly different (**Fig. 5R** marked as red ‘*’). Based on these analyses, TrimNN also found that motif ‘AEC’, a Homeostatic Interaction Motif, concatenating another cell type epithelial as a size-4 motif, was significant in disease and control. Similar analyses were performed to identify Shifted Interaction Motifs in **Supplementary Notes 3**. Using TrimNN, we recognized the decline of dominant CD4 T-cells in spatial space may be linked to colon carcinoma, where CD4 T-cells had more spatial relations with other cell types^46^.

## DISCUSSION

Spatial omics have significantly advanced our understanding of the nuanced cell organization within tissues at the cellular level. Complementary with top-down approaches such as clustering approaches, TrimNN enables the characterization of sizable CC motifs and provides a new bottom-up angle in spatial omics analysis. (1) It overcomes the limitation of clustering approaches. Without arbitrary parameters in clustering, the bottom-up approach identifies topological building blocks as countable CC motifs to represent CNs. (2) It provides intrinsic interpretability of topological building blocks of CNs. Easily interpretable biologically at the cellular and gene levels, and pathologically intertwined with clinical phenotypes, the results of TrimNN demonstrate the colocalization differences between repeated motifs and disease markers, which may correspond to different biological and pathological hypotheses. 3) It ensures better generalizability across samples. Without a complex machine learning model with uninterpretable embeddings dependent on trained samples, CC motifs as explicit interpretable representations are robust to differentiate CNs.

Mathematically, TrimNN offers an accurate, unbiased, efficient, and robust approach to quantifying CC motifs as interpretable building blocks of cell organization. The proposed work formulates the pattern quantification problem in counting subgraph occurrences, simplifying the task to a biologically constrained problem, which can be solved by a supervised topological representation learning framework using triangles as the inductive bias. TrimNN’s effectiveness depends on its ability to 1) simplify the NP-complete subgraph matching problem on the universal graph to a well-defined set of biologically meaningful triangulated graphs, 2) decompose the challenging isomorphic counting regression problem on the entire graph to many straightforward binary present/absent prediction problems on small graphs, which makes it possible to estimate biologically meaningful top overrepresented CC motifs with relative values, 3) using PE based graph representation methods to empower the expressive power of the model in spatial omics, and 4) using supervised learning approach to move the computational tasks to the training process to accelerate the inference to facilitate users’ practice. This work paves the way for disclosing the biological mechanisms underlying multicellular differentiation, development, and disease progression.

Biologically, TrimNN opens new opportunities for discovering complex mechanisms in complex diseases using spatial omics. The idea of CC motifs can be treated as an explicit representation that simplifies the cell organization and preserves the topological relations in the context of triangles from Delaunay triangulation, a fast and reliable mathematical process on spatial spaces as a CC. Using an interpretable LR model in the CRC study, the results of TrimNN robustly differentiate phenotypical and pathological characteristics of spatial omics samples. With TrimNN, we identified diverse CC motifs in spatial localization, cellular, gene, and pathway features in AD and colorectal carcinoma studies, corresponding to diverse biological and pathological mechanisms in complex diseases, which were often overlooked by conventional analysis.

Furthermore, finding CC motifs is also related to mining FTUs at the atlas level. For example, identifying tumor lysis syndrome^4^ (TLS) in cellular neighborhoods in cancer research helps to illustrate how the immune microenvironment plays a role in cancer progression. Using topology-based cell type combination in the spatial space, the abundance of CC motifs describes the dispersed and coherent cell organizations as shown in a demo system in **Supplementary Fig. 25**. In complex systems such as tumor microenvironment, these abundances can be used to describe the cell type boundary and gradient. Especially the abundance of specific cell types of tumor-immune mixing is also known to distinguish different organization scenarios such as a cold tumor, mixed tumor, and compartmentalized tumor, which are directly related to survival in triple-negative breast cancer^47^.

There are still some limitations of this work. Firstly, TrimNN enumerates the possible graph topologies and greedy search strategy to identify the large-size motifs, which has room for improvement with deep reinforcement learning approaches. Then, TrimNN is still estimated to reach the relative ranking in abundance and cannot be guaranteed to be exact. Furthermore, current settings in CCs may oversimplify the problem without any features on the edges and nodes. We may add essential features, e.g., morphology features, for specific applications. Future work will include analyzing large-scale spatial omics data to connect the idea of CC motifs and FTU, and assess the results in different categories of diseases from multiple independent data sources.

## METHODS

### Problem setting

Formally, we define the triangulated graph *G* as the CC inferred from spatial omics using Delaunay triangulation^13^, where *G* = {*V*, *E*} with |*V*| = *n* nodes and |*E*| edges. The size-*k* CC motifs is a subgraph with *k* nodes as *m_k_*, where *m_k_* is an induced subgraph of *G*. Here *m_k_* = (*V*′, *E*′) is defined as an induced subgraph if and only if when *V*′ ⊆ *V* and *E*′ = {(*u*, *v*) ∈ *E*|*u*, *v* ∈ *V*′}. *M_k_* is the set of all *m_k_* of size-*k*, and *M_k_* ⊆ *G*. The biological problem of identifying the overrepresented CC motifs can be modeled mathematically in finding the most overrepresented subgraph 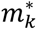 in *G*, where 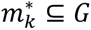 and 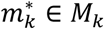. This challenge consists of a subgraph matching problem and a pattern growth problem built on it.

TrimNN aims to address the subgraph matching problem in the context of spatial omics, which seeks to define a function *F*(*G*, *m_k_*) ∈ ℕ, estimating the relative occurrence of the given *m_k_* in *G*. In our setting, this problem can be quasi-divided and conquered by summarization of many sub-TrimNN problems. The goal of sub-TrimNN is to build a reliable binary prediction model *f*(*g*, *m_k_*) ∈ [0,1], where 0 presents *m_k_* is absent in graph *g*, and 1 represents presence, *g* ⊆ *G*. With sub-TrimNN on enclosed graphs centered by each node, TrimNN is the summarization of results from all sub-TrimNN in the graph, as Eq. (1):

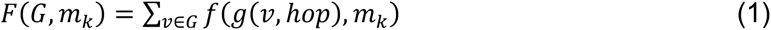

Where *g*(*v*, *hop*) is the enclosed graph as the neighborhoods of node *v* ∈ *V* with *hop* ∈ [1,2,3, …], and *g*(*v*, *hop*) ⊆ *G*. Generally, the value of *hop* is related to the length of the longest path of *m_k_*. Here, we use *hop* = 2 in all the analyses to make enclosed graphs *g* is in a similar size of *m_k_*.

We use a fast and reliable *F*(*G*, *m_k_*) from TrimNN to address the problem of pattern growth. Using searching processes, the final target is to get a top overrepresented set 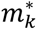 has the maximum relative abundance in Eq. (2):

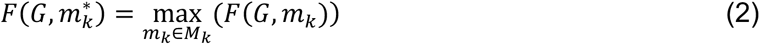

If both case and control samples are available, *F*′(*G_case_*, *G*_*control*_, *m_k_*) is defined to find CC motif *m_k_* that differentiates *G*_*case*_, and *G_control_* in Eq. (3), where *G_case_*, represents CC from case samples and *G*_control_ represents CC from control samples. *F*′ can be any function to describe the differences, including Fisher’s exact test or effective size. 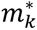 is the top overrepresented set mostly differentiated conditions in Eq. (4).

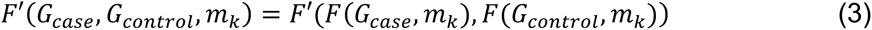

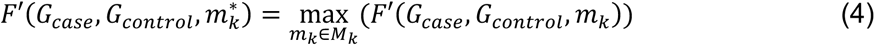

### TrimNN in subgraph matching

We decompose the regression problem of TrimNN in the CC into many binary classification problems in enclosed graphs centered by each node of the triangulated graph. This classification problem on each enclosed graph is solved by sub-TrimNN. The input of sub-TrimNN is a pair of subgraphs of query *m_k_* and the target triangulated graph *g*. To better represent the topological information of subgraphs and graphs, we use an empowered GNN based on GIN and a shortest distance positional encoding *PE*, which can be denoted as Eq. (5):

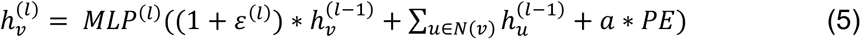

where 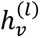 is the learned embedding of node *v* at the *l*-th layer. MLPs are multi-layer perceptrons, 𝜀 is a fixed scalar and *N*(*v*) is a set of nodes adjacent to *v*, and 𝑎 is the scaling factor for controlling the strength of *PE*. Here, *PE* as the Shortest Distance Positional Encoding^24^ is adapted to encode the relative positions of nodes in a graph based on the shortest path distances between them described as Eq. (6):

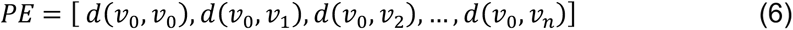

where 𝑑(*u*, *v*) denotes the shortest distance between two nodes *u* and *v*. For all nodes *v* in the graph *g*, we first selected one endpoint *v*_0_ of the shortest path in the whole graph *g* as the starting point. Then the shortest distance from this point was calculated for all other vertices.

After obtaining *PE* for the paired query *m_k_* and target *g*, we added these PE to the learned graph embedding from GIN. Then, learned node representations are passed through a graph max pooling layer^48^ to get the graph representations. After the sigmoid function activates linear layers, sub-TrimNN outputs the binary predictions as 1 as presence and 0 as absence. The whole training process aims at minimizing the cross-entropy loss function of known presence/absence relations as Eq. (7).

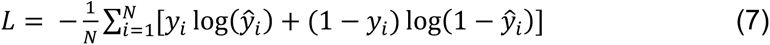

where *N* is the number of samples, 𝑦*_i_* is the true label for the 𝑖-th sample (0 or 1). 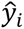 is the predicted probability for the 𝑖-th sample.

After trained sub-TrimNN *f*(*g*, *m_k_*), TrimNN estimates the abundances of *F*(*G*, *m_k_*) by summarizing sub-TrimNN predictions on each node’s enclosed graph as Eq. (1).

Theoretically, both the time and space complexity of sub-TrimNN inference are 𝑂(|*V*| + *k*), which is linear to the node sizes of the input subgraph and the triangulated graph. The time complexity of the entire TrimNN inference is the graph node size multiples the sub-TrimNN on all the enclosed graphs, which is 𝑂(|*V*| ∗ (*hop* ∗ *k* + *k*)), the time complexity of building an enclosed graph is 𝑂(|*V*|*^hop^*). The space complexity of the entire TrimNN inference is 𝑂(|*V*|*^hop^* ∗ (*hop* ∗ *k* + *k*)), where *hop* is defined in generating the enclosed graph. On the other hand, the space complexity of VF2 is of order 𝑂(*V*) and time complexity is 𝑂(*V*! ∗ *V*).

### Greedy strategy in pattern growth

In the pattern growth process, we use the function *F*(*G*, *m_k_*) from the subgraph matching in Eq. (2) to find 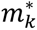 for a specified size *K* with a serial of greedy strategy. Starting from small sizes of CC motifs of size-*k*, where *k* = 1,2, *or* 3, we enumerate all possible subgraphs *m_k_*, and then obtain their corresponding predicted occurrence values using trained *F*(*G*, *m_k_*). For studies with case and control conditions, we calculate the *p*-value of Fisher’s exact test to identify the most significant CC motifs between different conditions. If no case-control information is available, we select the subgraph with the maximum relative abundance as 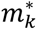. After obtaining 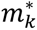 at size-*k*, we use it as a seed to enumerate all possible size-*k* + 1 subgraphs *M_k_*_+1_ based on 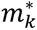

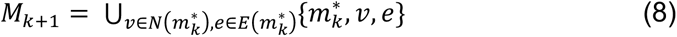

where 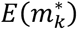 is a set of edges linked to the graph 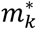. Each node type as a new node is selected with 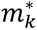 to get a new size-*k* + 1 graph. Similar to Eq. (1), 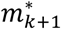 is defined as Eq. (9) in *M_k_*_+1._

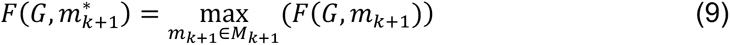

If both case and control samples are available, (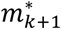)is defined as Eq. (10) in (*M*_*k* + 1_.

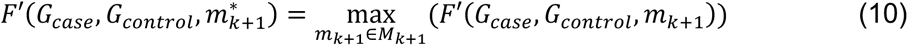

By iterating the process of Eq. (8) with incremental *k*, we can find 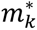 at specific size *K* in Eq. (9) or Eq. (10) if case and control samples are available, where *K* ∈ ℕ^+^.

### Constructing the training dataset

In spatial omics samples, the spatial relations between the cells can be modeled as a CC in a cell graph using the Delaunay triangulation^13^ on their spatial coordinates. In the generated cell graph composed of triangles after triangulation, each node denotes a cell and is labeled with a cell type, and each edge represents a hypothetical spatial relation between two cells.

We build a comprehensive synthetic training set with ground truth presence/absence relations between pairs of query CC motif (subgraph) and target triangulated graph. The classical tool VF2^21^ generated the ground truth occurrences by enumerating all the possibilities and guaranteeing the exact results with a substantial computational cost. To preserve biologically meaningful diversity, we simulated 7 distinct CC motifs from size 3 to size 9 in various topologies (**Supplementary** Fig. 26). Given the context of routine spatial omics in ROIs for each CC motif, we constructed the corresponding triangulated graphs with varying node types of 8, 16, and 32. To simulate different sizes of CC motifs corresponding to varying sizes of target graphs, we generated triangulated graphs of sizes 16 and 32 for size-3 to size-6 CC motifs, and triangulated graphs of sizes 32 and 64 for size-7 to size-9 CC motifs. Each pair of the query and the target has the same number of node types. To preserve the diversity, we generated 50 extended subgraphs with permutated node types for each CC motif. In total, we generated corresponding 1,000 triangulated graphs permutating node types for each extended subgraph. To ensure a balanced ratio of positive and negative relations in presence and absence, we controlled the proportion of positive to negative samples at 1:1 in data generation. We split and set the generated data into training, validation, and testing sets in a ratio of 8:1:1. Noteworthily, to fairly test model performance and objectively evaluate the model’s actual generalization ability, we constructed an independent test set. For each type of CC motif, we selected 50 entirely new permutations and generated 100 triangulated graphs for each permutation.

### Evaluation performance on synthetic data

We selected CC motif sizes ranging from size-3 to size-9 in the generated synthetic data to demonstrate the power of TrimNN with ground truth. As a binary prediction task of subgraph matching in the synthetic dataset, the performance was evaluated by precision, recall, F1 score, and MCC on the generated test set of varying sizes and node types. For NSIC regresses continuous occurrence, we treat NSIC’s prediction on 0 as the query absent in the target graph, and any value larger than 0 as present in the target graph. We omitted VF2 in the performance comparison, for it has already been used to generate the ground truth at a substantial computational cost.

To evaluate the quantitative performance in the synthetic dataset, we used metrics related to recommendation tasks, such as Root Mean Square Error (RMSE) and Mean Absolute Error (MAE), to quantify the accuracy of the predicted occurrences. We selected three distinct CC motifs, one in size-3, one in size-4, and one in size-5 (**Supplementary Data 2-3**). Then, we enumerated all the possibilities in both 8 cell types and 16 cell types with VF2 using significant computational resources. As we value the biologically meaningful top-overrepresented CC motifs in practice, we highlighted whether these methods can successfully identify the top 5 and top 10 overrepresented candidates as CC motifs.

In scalability analysis, the query subgraph contains 9 nodes, and both the subgraph and the triangulated graph have 32 node types. All the experiments were performed on a workstation equipped with an Intel Xeon Gold 6338 CPU with one NVIDIA A100 GPU and 80G RAM.

### Data preprocessing

#### Processing spatial proteomics in colorectal cancer study

The CRC study^13^ included in the analysis uses the spatial proteomics approach CODEX at the single cell resolution, which contains 140 tissue regions from 35 advanced-stage colorectal cancer (CRC) patients with 56 protein markers and 29 distinct cell types. This study consists of 17 patients labeled as “Crohn’s-like reaction” (CLR) and the remaining 18 patients as “diffuse inflammatory infiltration” (DII). Each patient has CODEX data with two spots, and each spot has two regions. Coordinates and cell type annotations of the cells were from the original publication. For all 140 samples, the summarized occurrence numbers of size 1-4 CC motifs were used in the classification tasks to predict their corresponding patients in either CLR or DII. Here we chose a unified number 29 as the number of features in size 2 to 4 CC motif analysis in alignment with the size-1 29 features as the total number of cell types. To get the top size-4 motifs, we performed TrimNN based on the top 29 size-3 CC motifs as triangles.

#### Processing spatial transcriptomics in Alzheimer’s disease study

In the AD datasets using STARmap PLUS spatial transcriptomics in this study^32^, 8 AD samples of mouse brain tissues – two replicates of an 8-month-old AD and control together with a 13-month-old AD and control were utilized for analysis. The original study provided each cell’s coordinates and their cell type annotation. The CC was built using Delaunay Triangulation, size 1-3 CC motifs were identified by enumeration, and the size-4 CC motif was inferred through TrimNN. In the downstream analysis on the cellular level, we only took the disease samples with amyloid-β regions and combined replicates 1 and 2. We categorized the samples’ regions containing specific CC motifs and their extended 3-hop as motif regions, and all the remaining regions were complementary non-motif regions. Then, we performed the cell-cell communication analysis using CellChat and DEG analysis using DESeq2. The spatial localization of amyloid-β and cells’ spatial coordinates were used to compute the shortest distance from amyloid-β to motifs.

#### Processing spatial proteomics in colorectal carcinoma study

The study^44^ uses MIBI-TOF with 36 antibodies on colorectal carcinoma, and it composes 58 ROIs within the spatial information and expression at the proteomics level. 40 ROIs were from two disease patients and 18 ROIs were from two control patients. Both NCEM and TrimNN were used to analyze these 58 ROIs.

### Noise simulation

To further validate the robustness of CC motifs, we applied three types of simulations based on STARmap PLUS data in the AD study^32^. These simulations manually added noise to mimic limitations arising from sequencing technologies and data processing in practical analysis, including i) cell missing, ii) cell coordinates shifting, and iii) cell type misclassification.

i. Cell missing targets to mimic the limited sequencing capacity to identify cells in the spatial omics samples. Proportions of cells in the input at rates of 0.01, 0.05, 0.1, 0.2, and 0.5 were randomly removed.
ii. Cell coordinates shifting aims to mimic errors in sequencing or shifting in sample preparation to identify the spatial localization of the cells. For proportions of cells at rates of 0.01, 0.05, 0.1, 0.2, and 0.5, their coordinates were shifted proportion of 0.01, 0.05, 0.1, 0.2, and 0.5 with their average distances to nearest neighboring cells in random directions.
iii. Cell type misclassification aims to mimic errors in annotating cell types from spatial omics samples, possibly due to insufficient cell type annotations. In the simulation, original cell types were randomly shuffled in cells at proportions of 0.01, 0.05, 0.1, 0.2, and 0.5.

To test the robustness of CC motifs in abundance ranking, all the simulations are randomly generated 100 times. Spearman correlation was adopted to compare the abundance ranks of all CC motifs between the original and the noisy datasets.

To test the generalizability of machine learning models, random cropping was proposed in addition to noises from cell missing, cell coordinates shifting, and cell type misclassification. Random cropping aims to simulate smaller ROIs due to limited sequencing capacity or incomplete sample preparation in that only a portion of the original tissue is captured. In this simulation, new patches as part of the original samples were randomly cropped with 0.1, 0.2, 0.3, 0.4, 0.5, 0.6, 0.7, 0.8, and 0.9 proportions of the original width and height. Random cropping was performed 10 times in each parameter setting. As part of the original samples, these newly generated small patches have identical phenotypes and survival of the original data. In generalization analysis, the machine learning model was trained on all the original CRC data and tested on the distorted samples.

### Statistics Summary

#### False discovery rate (FDR)

To assess the significance of CC motifs across samples, Fisher’s exact test and Chi-squared test were adapted in the study, and they were adjusted using Bonferroni correction and Benjamini Hochberg method.

#### Effect size

In large sample sizes, even small differences can become statistically significant (i.e., the *p*-value can be very small), which might not be practically significant. Therefore, we used the Cramér’s V effect size with a moderate effect size threshold of 0.21 in addition to the *p*-values to determine the significance of the Chi-squared tests^49^. For the hypothesis test that the occurrence of cell type A is independent of the occurrence of cell type B, the effect size of the chi-squared test is as follows Eq. (11):

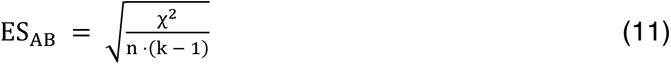

where χ^2^ is the chi-squared statistic, *n* is the total number of edges involved in each test, and *k* is 2 for the two-by-two contingency table. For the triangle of three types of cells in size-3 CC motifs, denoted as A, B, and C, we fixed one type of cell in the triangle and calculated the effect size for the test between two other types of cells as Eq. (12):

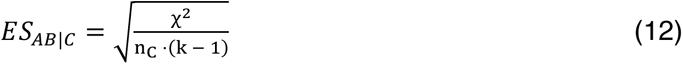

where *n_c_* denotes the total number of triangles that include at least one cell of C, and *k* is 2. The overall effect size was determined as the minimum value of *E*𝑆_*AB*|*C*_, *E*𝑆_*AC*|*B*_, and *E*𝑆_*BC*|*A*_, i.e., *E*𝑆_*ABC*_ = min (*E*𝑆_*AB*|*C*_, *E*𝑆_*AC*|*B*_, *E*𝑆_*BC*|*A*_), to ensure that the effect size of each chi-squared test is above the threshold.

#### Proportion test

We performed a two-sample, two-sided proportion test to test the significance between size-3 CC motifs and size-4 CC motifs. The null hypothesis (H₀) was that the proportions were equal between the two groups, i.e., H₀: the proportion of size-4 motifs within size-3 motifs is the same in both the disease and control groups.

### Parameter settings

#### Phenotypic classification in supervised learning

The top abundant CC motifs in sizes 1-4 were selected as the features to represent CNs. Specifically, the dense rank of the occurrence was scaled to [0, 1] by the function min_max. Classical machine learning models Logistics Regression, Radom Forest, and Support Vector Machine were adopted to perform classification tasks using 10 times 10-fold cross-validation following the same protocol as CytoCommunity. Comprehensive criteria such as F1 score, precision, recall, Matthew’s correlation coefficient (MCC), area under the precision curve, and area under the Receiver Operation Characteristics Curve (ROC-AUC) were used to measure the binary classification performances.

#### Survival curve

In survival analysis, we also summarized the overall occurrence of CC motifs through 140 regions for each size. The definition of whether a CC motif is enriched in a patient is by identifying whether it is among the top 29 motifs of its own sizes. Kaplan-Meier curves showing survival as a function of time for patients with and without CC motif enriched in cell organizations using R packages “*survival*” and “*survminer*”. The hazard ratio (HR) and *p*-value (P) were calculated using Cox regression analysis.

#### Cell-cell communication analysis

In the study of spatial transcriptomics datasets, CellChat^33^ performed ligand-receptor analysis using the ‘TruncatedMean’ method suggested by the official tutorial. In the cross-sample analysis, we retrieved the pathways related to targeted cell types of the single cell-cell interaction (targeted cell types as source or as target interacting with all other cell types) if the difference of cross-sample values is over 1. In the study of spatial proteomics datasets, we performed NCEM^45^ type-coupling and gene-wise analysis to supplement our size-2 triangulation analysis. All the parameters followed official tutorials.

#### Gene level analysis

We used package DESeq2^34^ to infer DEGs between motif and non-motif regions. Genes with *p*-values less than 0.05 were inferred as DEGs. GO and Pathway enrichment analysis were adopted by clusterProfiler^50^. Then, we applied the R package ‘*wilcoxauc*’ to perform the Wilcoxon Rank Sum test to identify marker genes in different motifs.

#### Spatial co-occurrence analysis

We used Squidpy^39^ function ‘*co_occurrence*‘ to calculate spatial co-occurrence probability between CC motifs and amyloid-β. The coordinates of CC motifs are the averaged coordinates of each node in the motif.

#### CytoCommunity and SpaceGM

We used CytoCommunity and SpaceGM to perform the supervised learning as the benchmark using the default parameter setting. Using the same protocol in CytoCommunity, we used the same 10 times 10-fold cross-validation and the same evaluation metrics for all the methods. It is notable that CytoCommunity performs training within in default 20 epochs, the output from the last epoch was shown as the final results.

## CODE AVAILABILITY

The source code of TrimNN is freely available at https://github.com/yuyang-0825/TrimNN.

## DATA AVAILABILITY

All relevant data supporting the key findings of this study are available within the article and its Supplementary Information files. The human CRC CODEX dataset used in the colorectal cancer case study is available at https://data.mendeley.com/datasets/mpjzbtfgfr/1. The STARmap PLUS sequencing data in Alzheimer’s disease case study are available at https://zenodo.org/records/7332091. The MIBI-TOF imaging data of colorectal carcinoma and healthy colon in spatial proteomics case study are available at https://doi.org/10.5281/zenodo.3951613.

## ACKNOWLEDGEMENTS

We thank Jiacheng Xie, Abdul Muqeeth for performing the analysis. This work is supported by National Institutes of Health grants R01DK138504 (to J.W. and Q.M.), R35GM126985 (to D.X.), R01GM152585 and P01AI177687 (to Q.M.), U19AG074879, R01AG019771, P30AG072976, U01AG072177, U01AG068057 (to M.Y.), NS121718 (to J.K.), R25HG012325 (to K.M.), the AnalytiXIN initiative (to J.W.), the Alzheimer’s Association grants AARF-22-722571 (to M.Y.), as well as the Pelotonia Institute of Immuno−Oncology (PIIO) (to Q.M.).

## AUTHOR CONTRIBUTIONS

Conceptualization: J.W., Q.M., and D.X.; methodology: J.W. and D.X.; software coding:

Y.Y. and S.W.; data collection and investigation: S.W., Y.Y., K.M.; data analysis: S.W., Y.Y., J.L., M.Y., K.M., J.K., A.M.; software testing and tutorial: Y.Y. and S.W.; manuscript writing, review, and editing: J.W., Y.Y., S.W., A.M., Q.M., and D.X.

## REFERENCES

1 Ruitenberg, M. J. & Nguyen, Q. H. Cellular neighborhood analysis in spatial omics reveals new tissue domains and cell subtypes. Nat Genet 56, 362–364 (2024). 10.1038/s41588-023-01646-x

2 Palla, G., Fischer, D. S., Regev, A. & Theis, F. J. Spatial components of molecular tissue biology. Nat Biotechnol 40, 308–318 (2022). 10.1038/s41587-021-01182-1

3 Gulati, G. S., D’Silva, J. P., Liu, Y., Wang, L. & Newman, A. M. Profiling cell identity and tissue architecture with single-cell and spatial transcriptomics. Nat Rev Mol Cell Biol (2024). 10.1038/s41580-024-00768-2

4 de Souza, N., Zhao, S. & Bodenmiller, B. Multiplex protein imaging in tumour biology. Nat Rev Cancer 24, 171–191 (2024). 10.1038/s41568-023-00657-4

5 Wu, Z. et al. Graph deep learning for the characterization of tumour microenvironments from spatial protein profiles in tissue specimens. Nat Biomed Eng 6, 1435–1448 (2022). 10.1038/s41551-022-00951-w

6 Hu, Y. et al. Unsupervised and supervised discovery of tissue cellular neighborhoods from cell phenotypes. Nat Methods 21, 267–278 (2024). 10.1038/s41592-023-02124-2

7 Varrone, M., Tavernari, D., Santamaria-Martinez, A., Walsh, L. A. & Ciriello, G. CellCharter reveals spatial cell niches associated with tissue remodeling and cell plasticity. Nat Genet 56, 74–84 (2024). 10.1038/s41588-023-01588-4

8 Singhal, V. et al. BANKSY unifies cell typing and tissue domain segmentation for scalable spatial omics data analysis. Nat Genet 56, 431–441 (2024). 10.1038/s41588-024-01664-3

9 Liang, Q., Huang, Y., He, S. & Chen, K. Pathway centric analysis for single-cell RNA-seq and spatial transcriptomics data with GSDensity. Nat Commun 14, 8416 (2023). 10.1038/s41467-023-44206-x

10 Schiebinger, G. et al. Optimal-Transport Analysis of Single-Cell Gene Expression Identifies Developmental Trajectories in Reprogramming. Cell 176, 928–943 e922 (2019). 10.1016/j.cell.2019.01.006

11 Bhate, S. S., Barlow, G. L., Schurch, C. M. & Nolan, G. P. Tissue schematics map the specialization of immune tissue motifs and their appropriation by tumors. Cell Syst 13, 109–130 e106 (2022). 10.1016/j.cels.2021.09.012

12 Jain, Y. et al. Segmenting functional tissue units across human organs using community-driven development of generalizable machine learning algorithms. Nat Commun 14, 4656 (2023). 10.1038/s41467-023-40291-0

13 Schurch, C. M. et al. Coordinated Cellular Neighborhoods Orchestrate Antitumoral Immunity at the Colorectal Cancer Invasive Front. Cell 182, 1341–1359 e1319 (2020). 10.1016/j.cell.2020.07.005

14 Lake, B. B. et al. An atlas of healthy and injured cell states and niches in the human kidney. Nature 619, 585–594 (2023). 10.1038/s41586-023-05769-3

15 Greenbaum, S. et al. A spatially resolved timeline of the human maternal-fetal interface. Nature 619, 595–605 (2023). 10.1038/s41586-023-06298-9

16 Wong, E., Baur, B., Quader, S. & Huang, C. H. Biological network motif detection: principles and practice. Brief Bioinform 13, 202–215 (2012). 10.1093/bib/bbr033

17 Patra, S. & Mohapatra, A. Review of tools and algorithms for network motif discovery in biological networks. IET Syst Biol 14, 171–189 (2020). 10.1049/iet-syb.2020.0004

18 Kashtan, N., Itzkovitz, S., Milo, R. & Alon, U. Eiicient sampling algorithm for estimating subgraph concentrations and detecting network motifs. Bioinformatics 20, 1746–1758 (2004). 10.1093/bioinformatics/bth163

19 Wernicke, S. & Rasche, F. FANMOD: a tool for fast network motif detection. Bioinformatics 22, 1152–1153 (2006). 10.1093/bioinformatics/btl038

20 Ullmann, J. R. An algorithm for subgraph isomorphism. Journal of the ACM (JACM) 23, 31–42 (1976).

21 Cordella, L. P., Foggia, P., Sansone, C. & Vento, M. A (sub) graph isomorphism algorithm for matching large graphs. IEEE transactions on pattern analysis and machine intelligence 26, 1367–1372 (2004).

22 Qi, M., Cao, T. T. & Tan, T. S. Computing 2D constrained delaunay triangulation using the GPU. IEEE Trans Vis Comput Graph 19, 736–748 (2013). 10.1109/TVCG.2012.307

23. 23 Xu, K., Hu, W., Leskovec, J. & Jegelka, S. How powerful are graph neural networks? *arXiv preprint arXiv:1810.00826* (2018).

24 Li, P., Wang, Y., Wang, H. & Leskovec, J. Distance encoding: Design provably more powerful neural networks for graph representation learning. Advances in Neural Information Processing Systems 33, 4465–4478 (2020).

25 Liu, X. et al. in Proceedings of the 26th ACM SIGKDD International Conference on Knowledge Discovery & Data Mining. 1959-1969.

26 Ying, C. et al. Do transformers really perform badly for graph representation?Advances in neural information processing systems 34, 28877–28888 (2021).

27 Chen, H., Covert, I. C., Lundberg, S. M. & Lee, S.-I. Algorithms to estimate Shapley value feature attributions. Nature Machine Intelligence 5, 590–601 (2023).

28 Liu, M. et al. The crosstalk between macrophages and cancer cells potentiates pancreatic cancer cachexia. Cancer Cell 42, 885–903 e884 (2024). 10.1016/j.ccell.2024.03.009

29 Laumont, C. M., Banville, A. C., Gilardi, M., Hollern, D. P. & Nelson, B. H. Tumour-infiltrating B cells: immunological mechanisms, clinical impact and therapeutic opportunities. Nat Rev Cancer 22, 414–430 (2022). 10.1038/s41568-022-00466-1

30 Prater, K. E. et al. Human microglia show unique transcriptional changes in Alzheimer’s disease. Nature Aging, 1–14 (2023).

31 Zhang, Y., Chen, H., Li, R., Sterling, K. & Song, W. Amyloid beta-based therapy for Alzheimer’s disease: challenges, successes and future. Signal Transduct Target Ther 8, 248 (2023). 10.1038/s41392-023-01484-7

32 Zeng, H. et al. Integrative in situ mapping of single-cell transcriptional states and tissue histopathology in a mouse model of Alzheimer’s disease. Nat Neurosci 26, 430–446 (2023). 10.1038/s41593-022-01251-x

33 Jin, S. et al. Inference and analysis of cell-cell communication using CellChat. Nat Commun 12, 1088 (2021). 10.1038/s41467-021-21246-9

34 Love, M. I., Huber, W. & Anders, S. Moderated estimation of fold change and dispersion for RNA-seq data with DESeq2. Genome Biol 15, 550 (2014). 10.1186/s13059-014-0550-8

35 Bellenguez, C. et al. New insights into the genetic etiology of Alzheimer’s disease and related dementias. Nat Genet 54, 412–436 (2022). 10.1038/s41588-022-01024-z

36 Ulland, T. K. & Colonna, M. TREM2 - a key player in microglial biology and Alzheimer disease. Nat Rev Neurol 14, 667–675 (2018). 10.1038/s41582-018-0072-1

37 Thambisetty, M. et al. Alzheimer risk variant CLU and brain function during aging. Biol Psychiatry 73, 399–405 (2013). 10.1016/j.biopsych.2012.05.026

38 Whyte, L. S. et al. Lysosomal gene Hexb displays haploinsuiiciency in a knock-in mouse model of Alzheimer’s disease. IBRO Neurosci Rep 12, 131–141 (2022). 10.1016/j.ibneur.2022.01.004

39 Palla, G. et al. Squidpy: a scalable framework for spatial omics analysis. Nat Methods 19, 171–178 (2022). 10.1038/s41592-021-01358-2

40 Kumari, S., Dhapola, R., Sharma, P., Singh, S. K. & Reddy, D. H. Implicative role of Cytokines in Neuroinflammation mediated AD and associated signaling pathways: Current Progress in molecular signaling and therapeutics. Ageing Res Rev, 102098 (2023). 10.1016/j.arr.2023.102098

41 Sobue, A., Komine, O. & Yamanaka, K. Neuroinflammation in Alzheimer’s disease: microglial signature and their relevance to disease. Inflamm Regen 43, 26 (2023). 10.1186/s41232-023-00277-3

42 d’Errico, P., et al. Microglia contribute to the propagation of Abeta into unaiected brain tissue. Nat Neurosci 25, 20–25 (2022). 10.1038/s41593-021-00951-0

43 Lei, X. et al. Immune cells within the tumor microenvironment: Biological functions and roles in cancer immunotherapy. Cancer Lett 470, 126–133 (2020). 10.1016/j.canlet.2019.11.009

44 Hartmann, F. J. et al. Single-cell metabolic profiling of human cytotoxic T cells. Nat Biotechnol 39, 186–197 (2021). 10.1038/s41587-020-0651-8

45 Fischer, D. S., Schaar, A. C. & Theis, F. J. Modeling intercellular communication in tissues using spatial graphs of cells. Nat Biotechnol 41, 332–336 (2023). 10.1038/s41587-022-01467-z

46 Valdeolivas, A. et al. Profiling the heterogeneity of colorectal cancer consensus molecular subtypes using spatial transcriptomics. NPJ Precis Oncol 8, 10 (2024). 10.1038/s41698-023-00488-4

47 Keren, L. et al. A Structured Tumor-Immune Microenvironment in Triple Negative Breast Cancer Revealed by Multiplexed Ion Beam Imaging. Cell 174, 1373–1387 e1319 (2018). 10.1016/j.cell.2018.08.039

48 Wang, Y. et al. Graph pooling in graph neural networks: methods and their applications in omics studies. Artificial Intelligence Review 57, 294 (2024).

49 Lin, M., Lucas Jr, H. C. & Shmueli, G. Research commentary—too big to fail: large samples and the p-value problem. Information systems research 24, 906–917 (2013).

50 Xu, S. et al. Using clusterProfiler to characterize multiomics data. Nat Protoc (2024). 10.1038/s41596-024-01020-z

